# Thermotaxis in an apolar, non-neuronal animal

**DOI:** 10.1101/2022.08.19.504474

**Authors:** Grace Zhong, Laurel Kroo, Manu Prakash

## Abstract

Neuronal circuits are hallmarks of complex decision making processes in the animal world. How animals without neurons process information and respond to environmental cues promises a new window into studying precursors of neuronal control and origin of the nervous system as we know it today. Robust decision making in animals, such as in chemotaxis or thermotaxis, often requires internal symmetry breaking (such as A-P axis) provided naturally by a given body plan of an animal. Here we report the discovery of robust thermotaxis behavior in *Trichoplax adhaerens*, an early-divergent, enigmatic animal with no anterior-posterior symmetry breaking (apolar), and no known neurons or muscles. We present the first quantitative and robust behavioral response assay in placozoan, which present an apolar flat geometry. By exposing *T. adhaerens* to a thermal gradient under a long-term imaging setup, we observe robust thermotaxis that occurs over timescale of hours, independent of any circadian rhythms. We quantify that *T. adhaerens* can detect thermal gradients of at least 0.1 ^*°*^C/cm. Positive thermotaxis is observed for a range of baseline temperatures from 17-22.5 ^*°*^C with motility trajectories to be well-described by a Lévy distribution. Interestingly, the organism does not maintain a fixed orientation while performing thermotaxis. Utilizing natural diversity in size of adult organisms (100um to a few mm), we also demonstrate a critical animal size above which thermotaxis behavior is hindered. Several TRP family homologs have been previously reported to be conserved in metazoans, including in *T. adhaerens*. We discover naringenin, a known TRPM3 antagonist, inhibits thermotaxis in *T. adhaerens*. The discovery of robust thermotaxis in *T. adhaerens* provides a tractable handle to interrogate information processing in a brainless animal. Understanding how divergent marine animals process thermal cues is also critical due to rapid temperature rise in our oceans.

## Introduction

It has been said that “nothing in neuroscience makes sense except in the light of behavior” ^1^ or “evolution”.^2^ How organisms integrate and transform sensory information into behavioral responses has both fascinated and confounded us since the days of Aristotle. Most behavioral responses in traditional biological model systems involve complex neuronal circuits.^3^ In these systems, complexity arises at emergent length scales – captured by the expression “more is different”.^4^ Reductionist approaches often fail to capture these systems completely. ^5^

Historically, “simpler” systems have been remarkably useful to identify key motifs in origins of behavior^6^ (which may be translatable to other organisms). A large number of such systems with fewer neurons (e.g. *C. elegans, Drosophila* larvae, lobster stomach) have provided hallmark studies enabling reduction of complex behaviors into sensory-motor transformations.^3,7,8^ However, even in these simpler model systems, it is still difficult to elucidate functional mapping at whole organism scale.^9^ In addition, biological systems usually demonstrate stereotypical architectures (for example, *C. elegans* always contains 302 neurons^9^) and hence do not allow an experimentalist to “scale the size” of the system in a meaningful way - making a comparative approach (so common in physics) almost impossible to implement. These challenges inspire us to study behavioral diversity in a broader range of non-model systems across the animal tree of life - with a possibility to circumnavigate these challenges. Since the explosion of multi-cellularity preceded the emergence of nervous systems, an illuminating question that arises is, how do multi-cellular animals without neurons or muscles, and without “nervous systems” in the current day definition, perceive and respond to environmental stimuli and make decisions? What are the limits of information processing capacity in these systems, and how do the limits map to the size of the organism? In our current work, we explore this space of sensory-motor transformation at “zero-brain limit”.

In the context of animal behavior, it is generally assumed that polarization of the system is a pre-requisite for generation of a directed migration in response to a stimulus.^10,11^ For most animals, this polarization/symmetry breaking is inherent in the body plan through anterior-posterior (AP) symmetry breaking.^12^ Even single cells such as neutrophils rapidly adopt a polarized morphology when presented with a chemical gradient.^13^ From single cells to cellular collectives, it is generally accepted that the direction of migration is governed by leading cells.^14^ For example, in the invasion of carcinoma cells, the leading cells are always fibroblasts.^15^ Some examples highlight the role of leader cell turnover, ^14^ such as in chemotaxing lymphoid cell clusters.^16^ Here we intend to explore if it is possible for an apolar animal without inherent AP symmetry breaking to make persistent long term decisions, such as in climbing up a shallow thermal gradient. What would govern the fundamental limits of sensitivity^17^ in such a system? How would these limits change depending on the size of the organism and total number of cells engaged in the process?

In order to better elucidate the role of size and symmetry breaking in taxis behavior, we seek to use a simple environmental cue as a handle. Temperature is a physiologically critical parameter that is crucial for survival of all life forms. ^18^ Thus temperature gradients are important ubiquitous environmental stimuli that influence behavior^6^ in both terrestrial and marine ecosystems. Furthermore, an urgency exists in understanding how marine animals (both adult and larval forms) process and adapt to thermal cues in face of rising ocean temperatures. Examples include settlement behavior of starfish^19^ and coral larvae,^20^ adaptation strategies of annelids in extreme temperatures,^21^ and worms that thrive near hydrothermal vents.^22^

Due to their broad applicability, thermal sensing capabilities and thermotaxis behavior have been reported ubiquitously in unicellular and multi-cellular systems. Most animals have been reported to be temperature-sensitive with a sensitivity on the order of 0.1^*°*^C/cm. Examples include humans (0.1^*°*^C) and human sperm (0.14^*°*^C/cm),^23,24^ *C. elegans* (0.1^*°*^C^25^) and *Drosophila* larvae (0.23^*°*^C/cm^7^). As an example of an extreme temperature-sensitivity, the nematode *Meloidogyne incognita* has a reported sensitivity limit below 0.001^*°*^C/cm.^26^ This example pushes the fundamental limits of what might be possible to build using molecular components in a thermally noisy environment. Focusing specifically in non-neuronal systems, thermotaxis behavior has been reported both in organisms that transiently form multicellular structures (*Dictyostelium discoideum* ^27^) and in single-celled organisms (phytoplanktonic flagellates ^28^ and *E. coli* ^29^).

In order to focus on how apolar non-neuronal animals might process environmental gradients, here we study the response of *T. adhaerens* to thermal gradients. *T. adhaerens* (Fig. 1a) belongs to phylum *Placozoa* and is the only known metazoan with no known anteriorposterior symmetry axis. The organism is composed of two epithelial layers in a flattened geometry - akin to a deflated beach ball. Furthermore, the animal does not have known neurons or muscles - it propels itself in an unusual fashion, coordinating millions of “walking cilia” ^30^ with asynchronous beats. This resulting “ciliary flocking” dynamics are rich yet descernable,^30–32^ providing a mechanical analog of behavioral control in these animals.^32^ Since both microscale dynamics and whole animal behavior can be studied in controlled conditions in the lab, *T. adhaerens* is a remarkable and experimentally tractable system to explore origins of information processing in non-neuronal animals. The animal further shows a natural yet large variation in size, allowing us to ask a fundamental question of how animal behavior varies with animal size.

**Figure 1.**
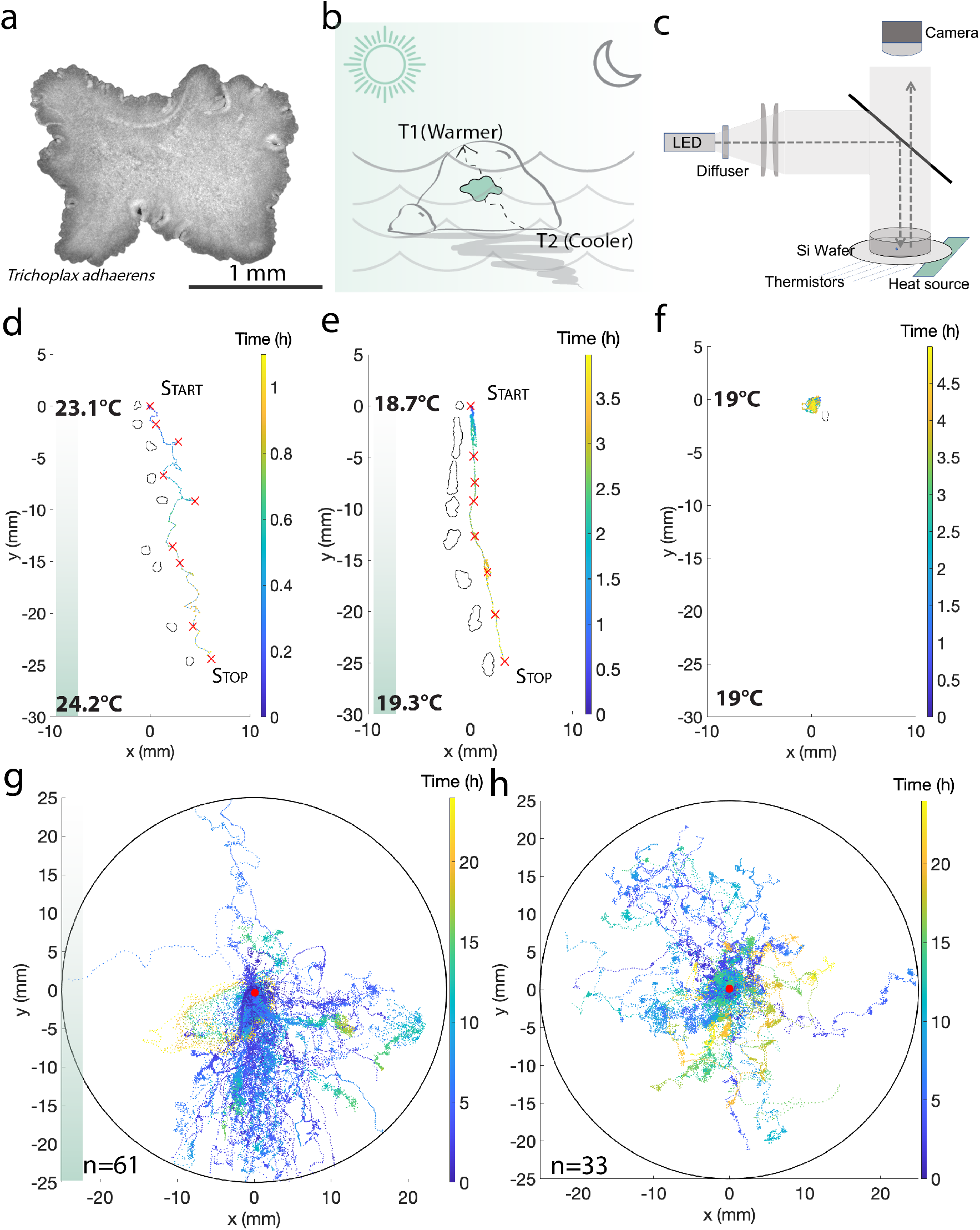
Discovery of robust thermotaxis in *T. adhaerens* - an organism in the phylum *Placozoa* (a)*T. adhaerens* has an apolar, flat geometry. (b) Thermal cues play an important role in their shallow-water natural environment. Temperatures would vary depending on factors like tides, seasons, time of day, and shadows. (c) Experiment setup using flexible heaters and overhead camera. (d-e) Sample organism trajectories show directed motion of organism towards higher temperatures under two different thermal gradients. Color bar indicates time. Segmented shape of the organism is shown at various time points. As is typical of *T. adhaerens* motile trajectories, the organism changes shape and projected two-dimensional area throughout the trajectory. (See SI Fig.2 for corresponding temperature profiles). (f) A sample control trajectory with no thermal gradient applied shows no directed motion. (g - h) Overlaid trajectories of biologically independent trials for heat (g, n=61) and control (h, n=33) conditions. T1 (temperature at warmer side of dish) range for trials shown is 17.3^*°*^C to 28.2^*°*^C, Δ*T* range is 0.008 to 0.42 ^*°*^C/cm.

Ecologically, *T. adhaerens* has been previously collected in tropical shallow waters around the globe, crawling on surfaces covered with algae mats. Large temperature fluctuations are common in shallow waters due to factors such as time of day, rock coverage, and seasonal changes in water temperature (Fig. 1b). Thus, being able to respond to temperature fluctuations can play an important role in the survival of any benthic species. Although the complete life cycle of any reported species in the phylum *Placozoa* (including *T. adhaerens*) has not been recapitulated in the lab, adult animal cultures can be grown at scale in lab conditions. This allows us to address behavioral questions in the context of this apolar, aneural system. Here we explore the “zero-brain limit” in animal behavior - a condition that can be reasonably linked to precursors of early animal forms.

Herein we report the discovery of robust thermotaxis in *T. adhaerens*. Positive thermotaxis is robustly observed across a wide temperature range. Intriguingly, thermotaxis behavior is observed over a period of several hours while the organism does not maintain a constant orientation with respect to the axis of the thermal gradient. Characterizing the trajectories using quantitative behavioral analysis approaches enables us to elucidate relevant timescales for integration of gradient information. We also find that the thermotaxis behavior can be well-described by a Lévy distribution. Further, we quantify the sensitivity of *T. adhaerens*’s ability to detect spatial thermal gradients to be least 0.1^*°*^C/cm. We further discover a non-monotonic relationship of thermotaxis effectiveness as a function of size. This provides a context of an optimal animal size range that is capable of reading thermal gradients and executing a taxis behavior. Finally, we lay the groundwork for identifying molecular underpinnings of this behavior by demonstrating reversible inhibition of thermotaxis by naringenin (a known TRPM3 antagonist), suggesting involvement of transient receptor potential (TRP) channel homologs known to exist in *T. adhaerens*. Thus, our work establishes a link between thermosensory circuitry and “ciliary flocking”.

## Results

### *T. adhaerens* demonstrates robust positive thermotaxis

To probe how *T. adhaerens* reacts to thermal gradients, we set up a 5.25cm diameter (approximately 3cm deep) experimental chamber with silicon wafer floor biased on one side with a polyimide flexible heater (Fig. 1c). Two independent methods were used for measurement of temperature gradients - thermistors lining the bottom of the silicon wafer for some chambers and an overhead infrared camera based setup for chambers without thermistors. Since organisms are cultured at approximately 18 ^*°*^C, we range our assay from 17.3^*°*^C to 28.2^*°*^C. As controls, and to ensure parameters other than the thermal gradient do not influence our assay, we performed the experiments at various times of day to rule out the effect of potential circadian cycles (SI Fig. 1a). Prior to running any experiments, organisms are starved for a few hours until they shed their undigested algae coat to ascertain that food stimuli and associated gradients are not present in the dish. Experiments are conducted sequentially with a single organism in the chamber at a time to eliminate potential effects of inter-organism communication. To avoid any influence of light, all experiments are conducted in the dark with only diffused overhead uniform illumination - just enough to spot the animal. See supplementary information for further details on controls experiments.

When thermal gradient is applied, *T. adhaerens* displays directed motion towards the warmer side of the arena (Fig.1d-e, Supplementary Video 1). In contrast, without an applied thermal gradient (Fig.1f, Supplementary Video 1), in a similar timeframe, the organism tarries near its starting point. We repeated the thermal gradient and control assays 61 and 33 times respectively. Comparing the motility tracks between these two conditions, it is evident that thermotaxis in *T*.*adhaerens* is a highly robust phenomenon (Figs. 1g-h). The application of a thermal gradient generates a very small surface flow of approximately 0.03mm/s (determined by tracking a floating debris). As an additional control to ensure that the observed behavior is not a result of this concomitant flow, we introduce a much stronger, externally generated, 3mm/s confounding flow using a stir bar. We show that the organism is still able to find the thermal gradient (SI Fig.1b, Supplementary Video 6), demonstrating that this phenomenon is unperturbed by these flows.

Next, we look more closely at thermotaxis motility trajectories using a range of relevant parameters. Under thermal gradients ranging from 0.008 to 0.42 ^*°*^C/cm, within the first 600 minutes, most organisms already show directed motion towards the thermal source (in the -y direction) (Fig.2a). In contrast, in the control condition, within the first 600 minutes, organisms tend to stay near the starting point or venture in random directions (Fig.2b). The distribution of momentary velocities for each condition reinforces the robustness of the phenomenon. When a thermal gradient is applied (Fig.2c), motion is clearly more directed towards the thermal gradient. This is in contrast with the even distribution of directions in the control condition (Fig.2d). When we introduce a sudden switch in gradient direction once the organism has already started moving up a thermal gradient, we observed that it takes the organism approximately 20 minutes to successfully reverse directions (Supplementary Video 4, Fig.2e).

**Figure 2.**
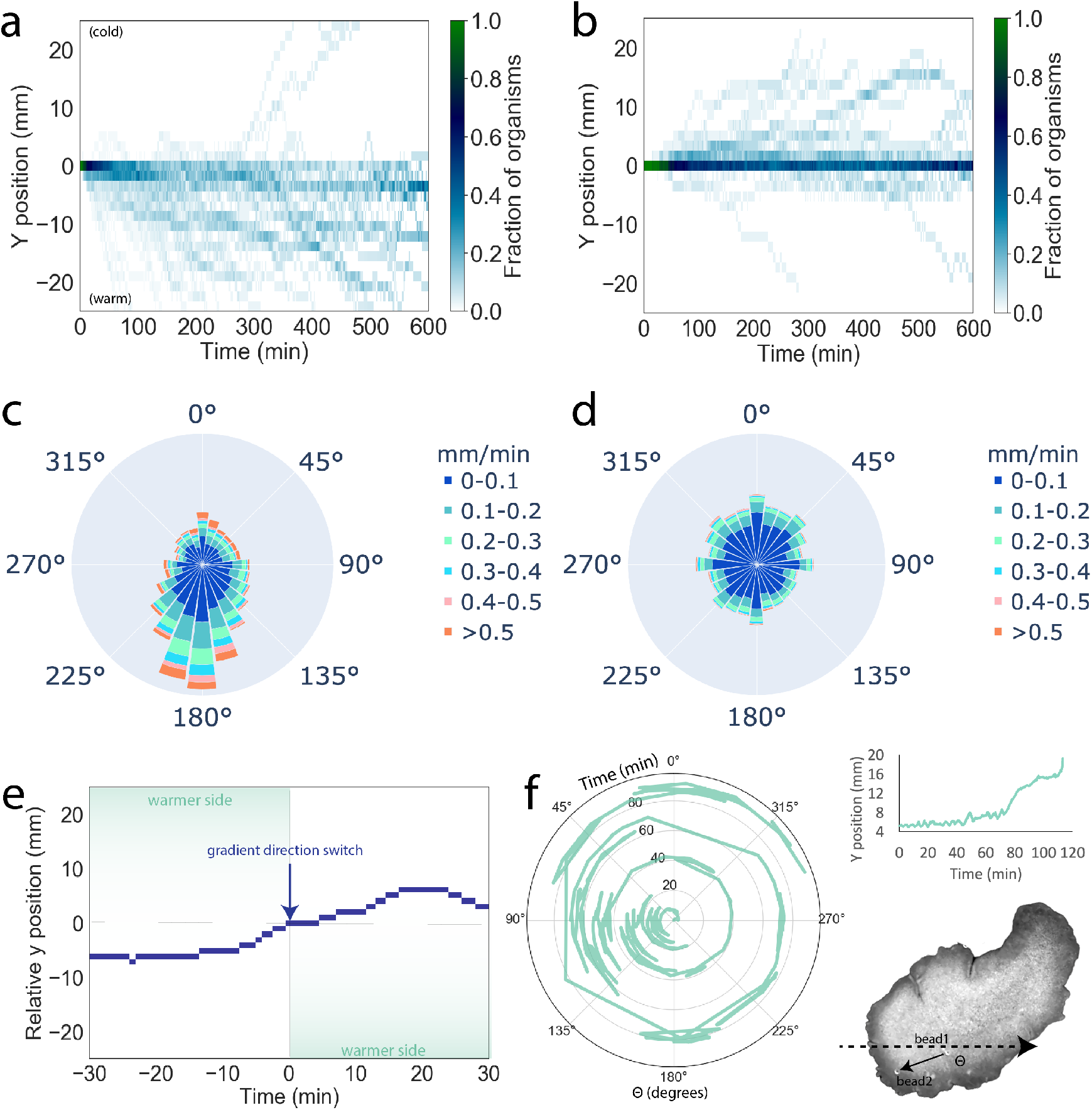
(a-b) Kymographs for heat (a) and control (b) conditions. When thermal gradient is applied, most organisms find the gradient within 10 hours. Without a thermal gradient, organisms still sometimes show directed motion, but the direction is often random, and it appears that more organisms tarry near the start point for longer before exploring. (c-d) Windrose velocity diagrams for heat (c) and control (d) conditions using overlapping window size of 10 min. When thermal gradient is applied, motion is directed towards the heat, and velocities parallel to the heat axis are also higher. Bars are grouped by velocity range and bar height normalized to total number of points. Radial axis limit is 0.1. (e) It takes approximately 20 minutes for the organism to switch directions when the direction of the thermal gradient is instantaneously flipped while the organism is already moving towards the warmer side (see Supplementary video 4). (f) During thermotaxis, *T. adhaerens* does not have a fixed orientation (see Supplementary Video 7). (Orientation is quantified as angle between “body axis” and positive x-axis. “Body axis” is defined using positions of two polystyrene beads attached to organism’s dorsal surface.)

### *T. adhaerens* does not maintain a fixed orientation with respect to thermal gradient axis while performing thermotaxis

Since all extant animals other than *Placozoa* break body axis symmetry via establishment of anterior-posterior symmetry, we explictly ask how the orientation of *Placozoa* - an apolar animal - changes during thermotaxis. To ask whether the organism polarizes to perform thermotaxis or remains apolar with varying orientation in the animal’s reference frame, we attach two WGA-coated polystyrene micro-beads (31.7 *μm* diameter) to the organism. We observe its thermotaxis behavior under higher magnification via tracking microscopy^33^ while also tracking the angle between the line segment connecting the two beads and the positive x-axis (Fig.2f, Supplementary Video 7). Notably, we find that *T. adhaerens* is able to establish a stable direction of movement along the thermal axis without a stable orientation in the frame of reference to the animal body (Fig.2f). This finding demonstrates a distinct set of processes must be at play in how thermosensory machinery is coupled to “ciliary flocking”^32^ in an ever-rotating frame of reference to enable this behavior.

A high degree of complexity exists across various length- and time-scales in *T. adhaerens* motility trajectories. In order to appreciate information embedded in these trajectories, it is useful to review how *T. adhaerens* walks on surfaces via rapidly beating cilia on it’s ventral surface. This unique mode of motility was recently characterized via direct real-time imaging of all cilia in a living motile organism. ^30–32^ Timescale of single cilia beat is *∼* 0.2 seconds, and timescale for locomotive dynamics of the organism is *∼*10’s seconds.^31^ Furthermore, something unique to *T. adhaerens* is the fact that its shape changes drastically while moving (Supplementary Video 7, Fig.1d-e). We recently established that the principles of *T. adhaerens* motility are embedded in the emergent phenomena of ciliary flocking dynamics.^30–32^ Interspersed in these trajectories, the organism also demonstrates short timescale events such as freezing (*∼*minutes), vortex insertion, and vortex ejection (which leads to transition from rotation to translation motion). These shorter timescale events then cascade up to the long timescale taxis which we observe, linking a vast range of length and time scales.

### Thermal gradient results in superdiffusive motion and increased long timescale correlation

Our thermotaxis assay gives us an organism-scale readout that embodies behavioral complexity across timeand sizescales. We next sought to quantitatively characterize these trajectories to find relevant metrics and timescales of the organism’s response to thermal gradients. On the longest timescale observable from our experiments, which is the “end-toend trajectory” timescale, the application of a thermal gradient manifests in a significant (p *<* 0.0001) increase in directionality index (Fig. 3a). This metric is also referred to sometimes as thermotaxis efficiency index.^34^ We also observe that with the application of a thermal gradient, overall trajectories are more aligned with the thermal gradient (SI Fig. 3b), and the organism spends more time in areas of the arena that are closer to the heat source than their starting point (SI Fig. 3a). On the shortest timescale of observation, we see a significant increase (p *<* 0.0001) in mean momentary velocity parallel to the thermal gradient (Fig. 3d). We confirm that this increase is not due to an increase in mean momentary speed (Fig. 3e, p*>*0.1). The slope of log-log MSD orthogonal to the thermal axis, as well as the orthogonal component of the momentary velocity, also do not differ significantly (p *>* 0.05) between the thermal gradient and control conditions (SI Fig.3c-d).

**Figure 3.**
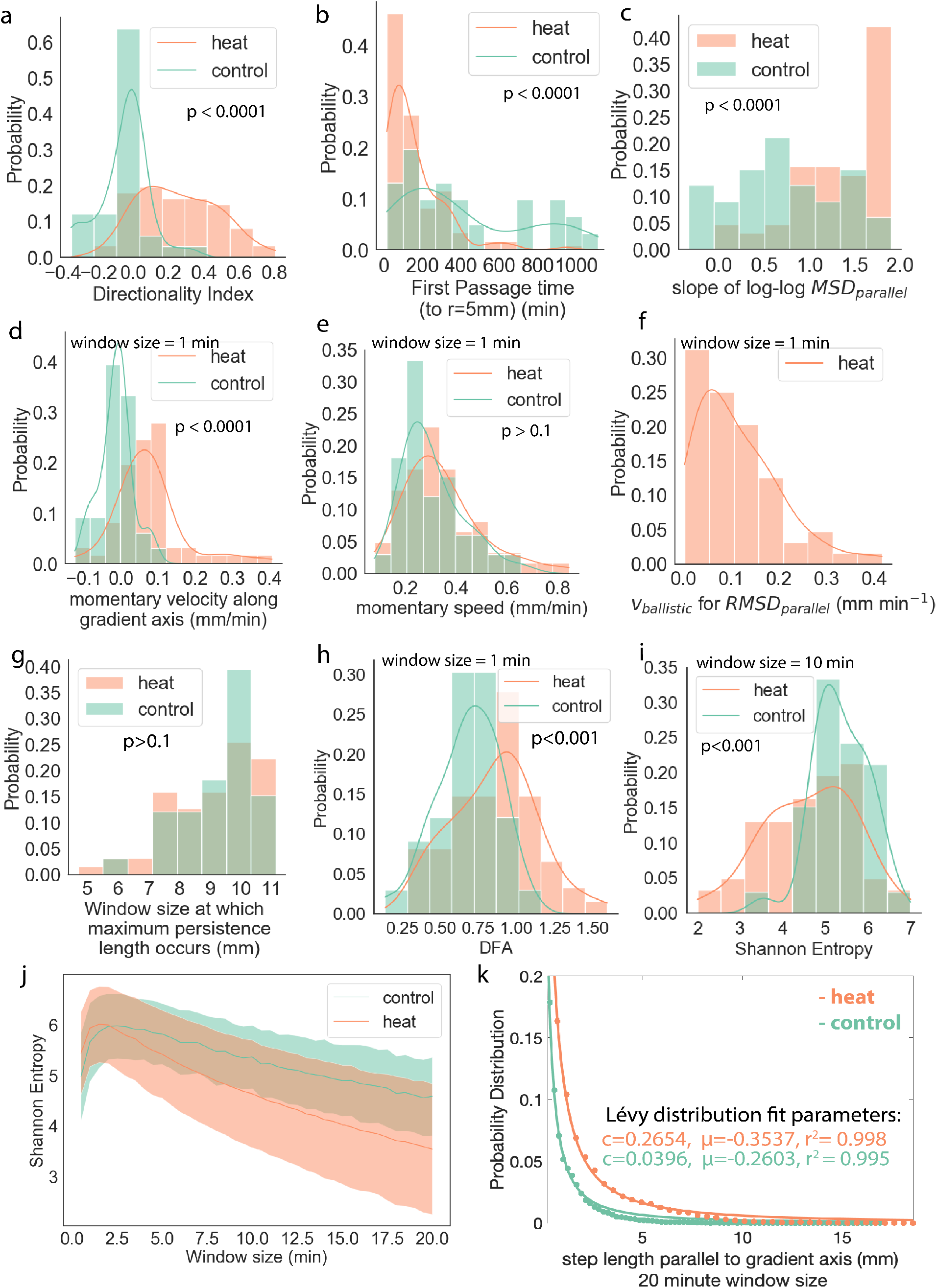
Various metrics to quantify directed motility in *T. adhaerens* undergoing positive thermotaxis. (a) With thermal gradient applied, we observe a significant increase in directionality index (overall displacement parallel to thermal gradient divided by total distance traveled). (b) We observe a significant decrease in first passage time (time for the organism to reach a radial distance of 5mm from its starting point) with application of thermal gradient. (c) Slope of log-log MSD parallel to thermal gradient indicates increase in directedness of motion. (d) Once the organism starts exploring, we observe an increase in mean momentary velocity parallel to the thermal gradient. (e) There is no significant increase in mean momentary speed (f) Once the organism starts exploring, the RMSD for trials under thermal gradient can mostly be fit using the linear ballistic motion estimate. (g) There is no significant difference between the window sizes that result in maximum persistence length for trials with and without thermal gradient. (h) Detrended fluctuation analysis shows a significant increase in long-range correlation in presence of thermal gradient. (i-j) Lower entropy seen with increasing sliding window size indicates increase in long-range correlation as a result of thermal gradient. (k) Step length distributions along gradient axis (for overlapping window size = 20 minutes) can be well-fitted by Lévy distributions.

We have previously established that time-scales of directional persistence is dictated by ciliary flocking dynamics, and in particular by the introduction and ejection of ciliary vortices.^32^ Next, we explore how the presence of a thermal gradient might affect persistence length in the organism trajectory. Looking at path persistence length,^35^ we find that window lengths of 7-11mm result in maximum persistence length, and there is no significant change with application of thermal gradient (Fig.3g, p*>*0.1). This suggests that the presence of a thermal gradient does not change inherent persistence patterns in organism-scale motion.

Even though the inherent persistence lengths in movement do not seem to be affected by the thermal gradient, the presence of the thermal gradient does increase long-timescale correlation. With an applied thermal gradient, we find the organism demonsrates superdiffusive motion when thermal gradient is applied (Fig. 3c). Under a thermal gradient, the root-mean-squared-displacement plot can be well-described by ballistic motion estimate (Fig.3f). The increase in long-range correlation with application of thermal gradient also manifests in significant differences between thermal gradient and control conditions in detrended fluctuation analysis (DFA), and significant decrease in entropy as we increase sliding window size (Fig. 3h-j). Thus, even though in both conditions we see “segments” of directed motion, the segments are more directionally aligned when a thermal gradient is applied.

### Motion along gradient axis can be described by Lévy distribution

The observation of segments of directed motion leads us to explore whether the trajectories can be described by known search models with heavy tailed step size distributions. We find that the step length distribution along the thermal gradient axis for our assays can be well-described by the Lévy distribution^36^ (Fig. 3k):

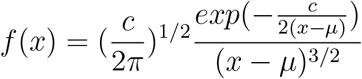

with *c* = 0.2654, *μ* = *−*0.35 and *c* = 0.0396, *μ* = *−*0.26 respectively for thermal gradient and control conditions, determined using a window size of 20 minutes and least squares regression fitting. Lévy walks have been previously used to describe search patterns in various taxa, including in dinoflagellate prey search^37^ and in individual swarming bacteria.^38^ The heavy tail distribution fit supports our observation of high propensity for directed motion (observed over tens of minutes) along gradient axis in presence of thermal gradient.

### *T. adhaerens* does not show preference for culture temperature

In addition to search strategy, thermotaxis behavior is also interesting in the context of ecology, adaptation, and size-scaling. So, next we ask how parameters such as absolute temperature, gradient strength, and size of organism affect thermotaxis behavior. We explore if, like for other organisms such as *C. elegans, T. adhaerens* thermotaxis shows a preference for culture temperature. We test T1 ranging from 17 ^*°*^C to 22.5 ^*°*^C, which spans both sides of our culture temperature (18 ^*°*^C). *T. adhaerens* does not appear to show a preference towards its culture temperature. Within the limited range of temperatures tested, there is no absolute-temperature dependent trends in directionality index (Fig.4a), and the organism shows robust positive thermotaxis through this entire range of T1.

**Figure 4.**
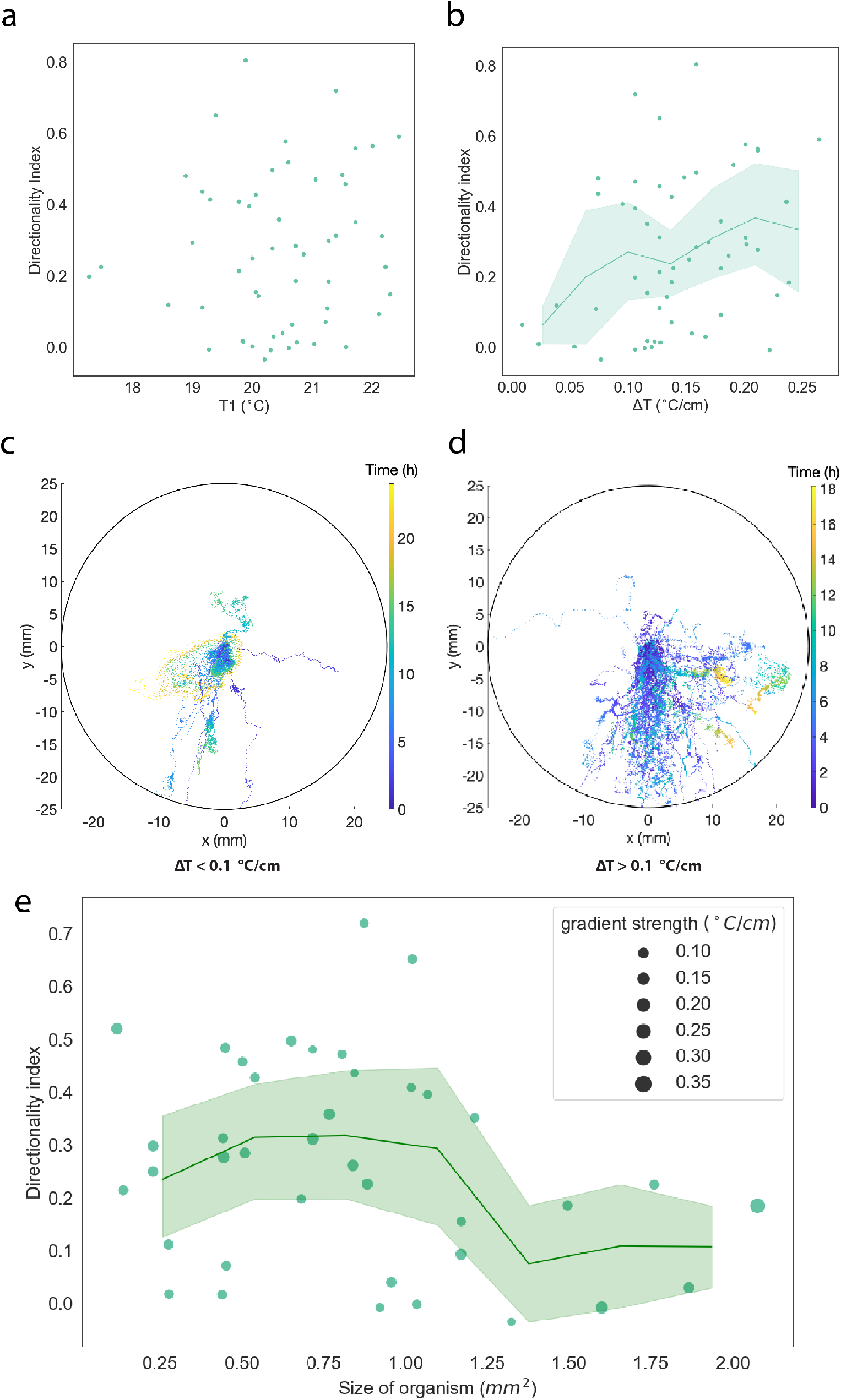
*T. adhaerens* does not show a preference for culture temperature, but thermotaxis behavior may be gradient strength and size-dependent. (a) There is no apparent correlation between T1 (temperature at warmer side of the arena) and directionality index for T1 spanning both sides of culture temperature (18 ^*°*^C). (b) There appears to be a weak correlation between gradient strength and directionality index. (c-d) Binning trajectories by whether the gradient strength is below (c) or above (d) 0.1 °C/cm, we qualitatively see the organism may be less effective at finding or moving towards the gradient at low gradient strengths. However, even at low gradient strengths, the organism is mostly able to detect the gradient. (See SI Fig.4 for the trajectories for trials with the three lowest Δ*T* that we tried, see SI Video 2 for video of trial with shallowest gradient) (d) We observe a drop in directionality index in organisms larger than 1.25 *mm*^2^.

### *T. adhaerens* can detect gradients of at least of 0.1 ^*°*^C/cm

We next sought to determine whether the strength of the thermal gradient affects *T. adhaerens* thermotaxis. There appears to be a weak dependence of directionality index on Δ*T* (Fig. 4b), with higher Δ*T* corresponding to higher directionality index, albeit with fluctuation. Typically, a directionality index above 0 is considered positive thermotaxis. If we take 0 as the cutoff for positive thermotaxis, then 78 percent and 91 percent of organisms demonstrate positive thermotaxis at gradients below and above 0.1 ^*°*^C/cm, respectively. If we define “effective thermotaxis” as thermotaxis with a directionality index higher than 0.1, then 56 percent of organisms in our experiments effectively thermotax at gradient strengths below 0.1 ^*°*^C/cm (Fig.4c). At gradient strengths above 0.1 ^*°*^C/cm, 76 percent of organisms effectively thermotax. Thus, we can safely conclude that the organism is able to detect temperature gradients of at least 0.1 ^*°*^C/cm. The lowest temperature gradient that *T. adhaerens* is able to detect in our experiments was 0.008 ^*°*^C/cm (Supplementary Video 2, SI Fig.4).

### Effectiveness of thermotaxis may be lower above a certain organism size

*T. adhaerens* naturally shows orders of magnitude variations in size. To probe the effect of size on thermotaxis behavior, we select naturally occurring organisms of various sizes. In our experiments, the smallest organism measures 0.12 mm^2^ while the largest organism is 2.1 mm^2^ in size. Roughly estimating the area of each cell to be 30*μ*m^2^ and the organism as having two cell layers,^39^ this translates to a size range of less than 10000 cells to around 135000 cells.

Though the standard deviation is high, we observe a critical size (approximately 1.25mm^2^ from Fig.4d) above which thermotaxis is less efficient. Why there could be an optimal size range for stimulus-response leads us to ask what the thermosensory circuitry looks like. To get a handle on the circuitry, we ask what thermosensitive ion channels might be present in *T. adhaerens*.

### Transient receptor potential (TRP) channel homologs may be temperature sensors in *T. adhaerens*

In seeking to identify potential sensors mediating the thermotaxis response, we start by looking to transient receptor potential (TRP) channels. TRP channels have diverse known functions; they act as ubiquitous sensors in vertebrates and invertebrates alike and are responsive to numerous chemical and physical stimuli, including temperature.^40–44^ These channels are additionally interesting because they are often polymodally activated and are considered an example of signal integration at the level of the sensor itself.^44,45^

TRP channel homologs have been identified in the ocean. For instance, TRPA, TRPC, TRPM, TRPML, and TRPV channels were identified in the marine unicellular choanoflagellate *M. brevicollis*, demonstrating the evolution of these TRP subfamilies prior to the emergence of multicellularity. ^46^ TRPA and TRPML channels have also been identified from the genome of the sponge *A. queenslandica*.^46^ These channels might be involved in facilitating the (non-neuronal) responses observed in these organisms.

TRP channel homologs have also been identified from the *T. adhaerens* genome – specifically TRPM1-4, TRPML, TRPP1-2, and TRPV1-2.^46,47^ Of these potential TRP channel homologs identified, the mammalian counterparts for TRPV1-2^48^ and TRPM2-4^49–51^ have been previously reported to be temperature-sensitive (SI Table 1). The percent similarities of these hypothetical proteins with the canonical homologs in *Mus musculus* are not very high (SI Table 1), but when we did a blastp search using hypothetical *T. adhaerens* proteins, entries with highest percent similarity for hypothetical *T. adhaerens* TRPM3 and TRPM4 homologs were from the ocean – in particular, from corals (*Acropora millepora, Pocillopora damicornis*) for TRPM3 and from sea stars (*Patiria miniata, Asterias rubens*) for TRPM4 (Table 1). So for this study, we first focus on TRPM3 and TRPM4.

**Table 1:** blastp results with highest percent similarity to amino acid sequences of predicted *T. adhaerens* TRP channel homologs whose mammalian counterparts have previously been shown to be temperature-sensitive.

| *T. adhaerens* TRP channel homologs<br>(protein database accession) | blastp results with highest % similarity | % Similarity |
| --- | --- | --- |
| TRPM2 (XP_002114984) | XP_034144252 (*Esox lucius*) | 29.6% |
|  | XP_025915155 (*Apteryx rowi*) | 28.2% |
| TRPM3 (XP_002114617) | XP_044172604 (*Acropora millepora*) | 40.7% |
|  | XP_027040503 (*Pocillopora damicornis*) | 39.2% |
|  | CAC9446695 (*Owenia fusiformis*) | 37.4% |
| TRPM4 (XP_002116621) | XP_038079149 (*Patiria miniata*) | 39.6% |
|  | XP_033624225 (*Asterias rubens*) | 39.3% |
| TRPV1 (XP_002114713) | XP_035674128 (*Branchiostoma floridae*) | 39.8% |
|  | XP_022322068 (*Crassostrea virginica*) | 39.7% |
| TRPV2 (XP_002118208) | PVD25324 (*Pomacea canaliculata*) | 43.2% |
|  | XP_006825516 (*Saccoglossus kowalevskii*) | 41.3% |

In order to further explore the role of TRP channel homologs in *T. adhaerens*, we start with several drug perturbation assays. Thermotaxis behavior is interrupted by addition of 40 *μ*M naringenin, a known inhibitor of TRPM3^52,53^ (Fig. 5a, Supplementary Video 3). This inhibition of thermotaxis by naringenin is robust across 14 biologically independent trials (Fig.5b). The inhibition is temporary and reversible: most organisms recover positive thermotaxis behavior within one hour post-naringenin addition or following washout (Fig.5b,SI Fig.1d). The inhibition also occurs without concomitantly inhibiting organism motility (Fig.5c-d). Flufenamic acid (FFA), which is a reported inhibitor of the TRPM4 channel,^54^ also seems to inhibit thermotaxis (Fig.5c). However, FFA addition seems to also reduce motility (Fig.5d). In addition, FFA is known to inhibit a wide range of ion channels,^54^ so it is inconclusive whether the observed behavioral inhibition is due to sensory inhibition.

**Figure 5.**
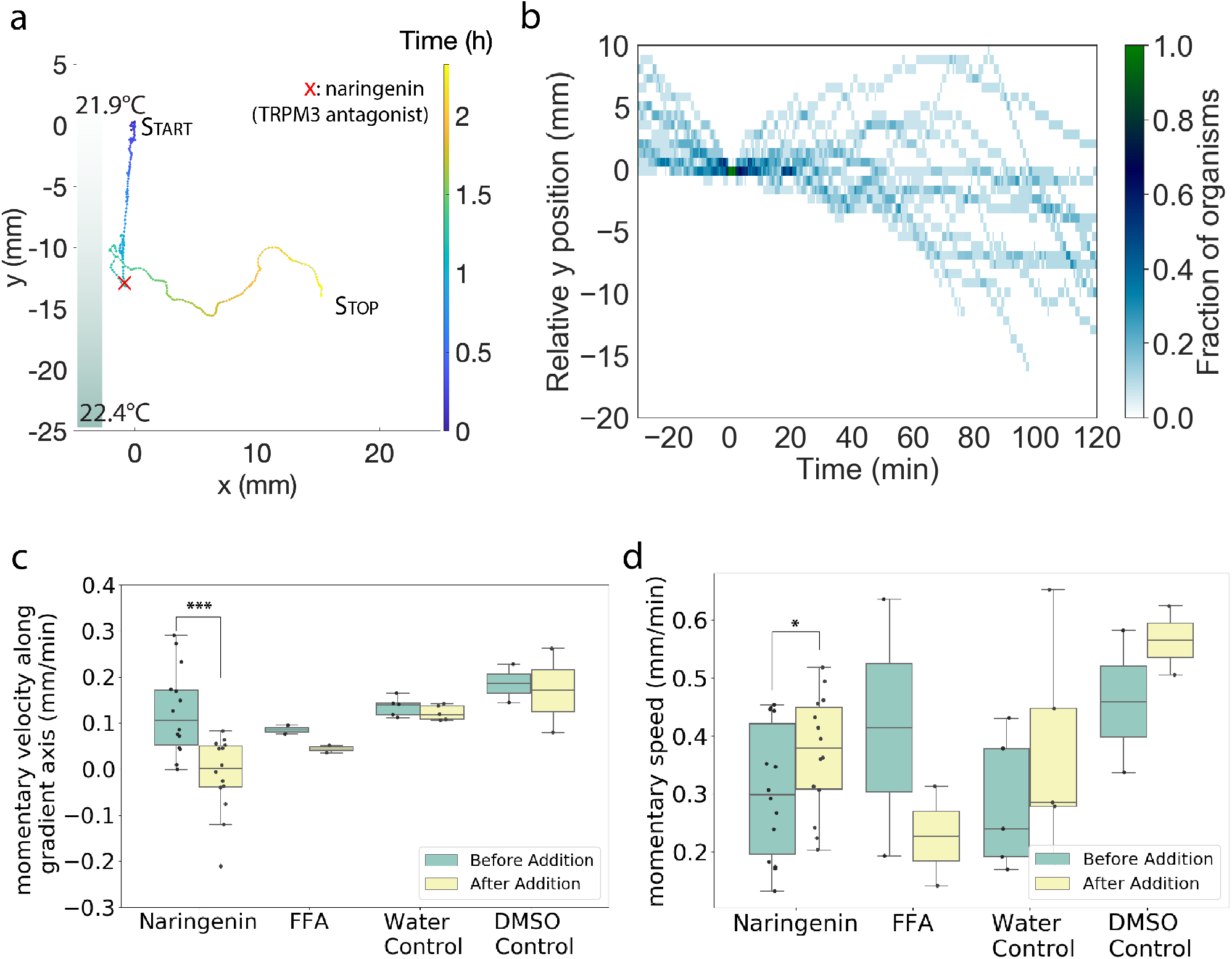
TRP channel homologs are potentially involved in signal cascade driving response to thermal stimulus. (a) A sample trajectory under the indicated thermal gradient, with 40 *μ*M naringenin (known TRPM3 antagonist) added at the time point indicated by the red x. Naringenin addition inhibits thermotaxis behavior, but the organism seems to recover after around an hour. (b) Kymograph showing positions along thermal axis relative to the position at naringenin addition for n=14. Naringenin robustly inhibits thermotaxis, and the inhibition appears to be transient, with most organisms recovering after an hour. (c-d) Boxplots showing mean momentary velocity along gradient axis (c) and mean momentary speed (d) in 30 minute time windows before and after addition of naringenin (TRPM3 antagonist) and FFA (TRPM4 antagonist). Sliding window size is 30s, non-overlapping windows. It appears both drugs inhibit thermotaxis (c), and the observed thermotaxis inhibition is not due to the DMSO solvent used nor the seawater that’s used for dilution (More details on controls in SI Fig.1c-d). Naringenin inhibits thermotaxis without reducing motility (interestingly, it leads to a slight increase in motility) while FFA appears to also reduce motility (d). *: p*<*0.05, ***:p*<*0.001

Thus, our data shows inhibition of thermotaxis by naringenin, which occurs without motility inhibition. This suggests that a TRP channel homolog (perhaps a TRPM3 channel homolog) may be a temperature sensor in *T. adhaerens*.

## Discussion

All benthic organisms face a complex seascape of environmental conditions. Many organisms such as slime molds and *C. elegans* show preference for their culture temperature, and their thermotaxis behavior can be altered by changing culture conditions. By running a range of experiments at various baseline temperatures, we find *T. adhaerens* thermotaxis does not exhibit a preference for culture temperature (18 ^*°*^ C). Thus, decision making might rely on a broad range of signals instead of one single preferred temperature. In the future, we intend to explore multi-sensory stimuli in our arenas. Decision making in a multi-sensory stimuli arena might also benefit from the fact that *T. adhaerens* does not break symmetry and remains apolar in its frame of reference.

Ciliary dynamics play a key role in understanding how *T. adhaerens* performs thermotaxis as an un-polarized tissue. To the best of our knowledge, *T. adhaerens* is the only organism that performs thermotaxis without polarization along the axis of the gradient as a prerequisite. Here we have an organism that does not have an inherent AP axis hence ideas like heading do not really apply. Rather, the motion and directedness are cascaded up from and governed by ciliary flocking dynamics, where coherent, directed motion is produced as a result of cell-cell interactions.^31^ By not aligning itself to a gradient for prolonged period of time (encode memory of gradient axis), sensory-motor systems might also be more adaptive in multi-sensory and variable environments.

We have experimentally demonstrated that there is an optimal organism size range for effective thermotaxis. Beyond size, anomalous perimeter to area ratio also hinders successful thermotaxis (as in Supplementary Video 5). These findings lead to many interesting questions in the realm of sensory-motor systems. Previously, optimal cluster size in collective migration has been seen both theoretically^55^ and experimentally (in context of border cells in *Drosophila* egg chamber).^56^ Although these previous findings are relevant, significant differences exist in how *T. adhaerens* glides on surfaces due to coherent “ciliary flocking”. Since the numbers of cells and cilia in an organism scale linearly with size, above a certain size, coherence in ciliary flocks decreases with size, ^30^ making it more difficult for larger tissues to change course. We hypothesize that the behavior is sensor-limited at smaller sizes, while at larger sizes the behavior is coordination-limited.

In many sensory-motor systems, it’s possible to ask what an optimal sensor-actuator ratio might be. Where might these sensors be located? Several models of integration of information along the sensory-motor axis are feasible in the frameworks of the discovery of thermotaxis described in *T. adhaerens*. Assuming that individual cilia themselves are sensory organelles, several models can be built for modulation of gliding motility as a function of thermal gradient.^32^ Such a possibility has been previously shown for light driven adhesion modulation in cilia in *Chlamydomonas*.^57^ In this case, cilia as sensors will scale lineraly with the size of the organism. Another possibility is existence of specific TRPM3 expressing sensory cells that upon activation release particular peptides or small molecules involved in *T. adhaerens* behavior modulation. We currently do not know where TRPM3 expressing cells might be located in the organism. Support for TRPM3’s role in driving peptide or small molecule release has previously been shown.^58,59^ Since several neuropeptides that modulate *T. adhaerens* have already been demonstrated,^60^ in the future we intend to explore how thermotaxis links to neuropeptide based behavioral control. In addition, it is also feasible that mucus trails observed previously^61^ in *T. adhaerens* might be able to encode spatial memory and modulate thermotaxis.

Finally, since complex ciliary flocking dynamics such as vortex injection, ejection, merging and splitting are all involved in encoding motility behavior in *T. adhaerens*, a higher order combination and control of these phenotypes might be involved in encoding thermotaxis. As we demonstrate in our experiments, organisms take hours to detect and respond to thermal cues. The vast gap in time scales between ciliary flocking dynamics and the time for detecting and climbing a gradient presents a fertile ground to explore a number of ideas from sensor to motor transduction.

Our current work in this non-model system - *T. adhaerens* - lays the foundation for studying higher order sensory and information processing capabilities of this seemingly simple animal. By building a rigorous experimental framework of tracking animals making decisions in complex artificial sensory seascapes, we believe a large number of questions pertaining to information processing in early multi-cellular life forms can be now formulated. For instance, how localized is the processing of information such as reading a complex gradient?

## Conclusion

In this work we show the discovery of robust thermotaxis in *T. adhaerens*, a simple multicellular organism of the phylum *Placozoa* with no known neurons or anterior-posterior symmetry breaking. We develop a framework for conducting systematic, long-term behavioral assays for *T. adhaerens*. To the best of our knowledge, this is the first quantitative and rigorous behavioral assay for this non-model organism. We characterize the trajectories using a myriad of metrics relating to directedness of motion and physics of behavior, and find that the trajectories can be well-described by a Lévy distribution. Further, we find that *T. adhaerens* can detect thermal gradients of at least 0.1^*°*^C/cm. Notably, we show that unlike many other organisms that perform thermotaxis, *T. adhaerens* does not show a temperature preference that is related to culture conditions. It also does not maintain a fixed orientation while performing thermotaxis. We further report the potential presence of an optimal size range for effective thermotaxis. Combined with our finding that TRP channel homologs might be potential temperature sensors in this system, we establish thermotaxis in *T. adhaerens* as a tractable behavioral handle to ask fascinating questions surrounding how an apolar tissue can respond to sensory signals through distributed signal processing. The rigorous behavior assay we describe and our findings create a path to explore decision making mechanisms that control both short- and long-term behavior in this non-model, multi-cellular system at the “no brain limit”.

The question of different types of strategies in responding to thermal cues and what they could mean in an ecological context could also take on a particular importance in the context of climate change - how might changing climate affect behavioral response, and what impact could it have on organism survival? This question carries a somber significance for an organism where we have only discovered a handful of known species (less than 10 - all found in tropical waters) associated with the entire phyla.

## Methods

### Culture of *T. adhaerens*

Our cultures of *T. adhaerens* are propagated from original cultures that were a gift from L. Buss, Yale University. They are maintained in artificial seawater (ASW) with 3 percent salinity, in glass petri dishes. Cultures are kept on shelves in a dedicated room with temperature maintained at 18 ^*°*^C and light cycle of 18h light - 6h dark.

ASW is prepared to reach 3 percent salinity by adding Kent Marine Reef Salt Mix or CoralPro Salt Mix in Millipore water on a magnetic stirrer, and salt concentration is measured by ATC salinity refractometer.

The food source for the *T. adhaerens* culture is marine alga *Rhodomonas lens* (R. Lens), propagated from cultures that were gifted by C. Lowe lab (Stanford). R. Lens cultures are maintained in 500-mL sterile Erlenmeyer flasks either in dedicated incubators maintained at 14 ^*°*^C or in the same room as *T. adhaerens* (maintained at 18 ^*°*^C) on separate shelves. Micro Algae Grow from Florida Aqua Farms is used as a source of nutrients for R. Lens cultures. Algae flasks are shaken daily.

For our *T. adhaerens* cultures, we start by adding 250 *μ*L of filter-sterilized Micro Algae Grow per 1000 mL of filtered ASW to create an ASW stock. Then we fill the glass petri dish with 110mL of the ASW stock and 15mL of R. Lens. We let the plate rest on the shelf for 2-3 days to allow for a growth of an “algal mat” at the dish bottom, then we add 10-20 *T. adhaerens* to the plate.

We do a 30mL water refresh once or twice weekly, and also add 5mL of algae during plate refreshes if the organisms no longer look pink in color.

### Behavioral assays

*T. adhaerens* were placed in nutrient-free ASW for 4-8 hours prior to behavioral experiments in order for them to shed their undigested algae coat so as to remove chemical information from algae to isolate the effect of thermal information. They are then pipetted into the middle of the arena, which is filled with 25mL of nutrient-free ASW, the heat source is turned on, and the acquisition (using custom Python script), is started.

The chamber for all described behavioral experiments except for the orientation experiment is made from an approximately 3cm tall section of clear PVC piping with 2.06 inch inner diameter (McMaster Carr) attached to a silicon wafer (University wafers) with silicone sealant. To the back of the wafer is attached, with thermally conductive tape, either a flexible polyimide heating strip (Omega) or two 11W 5V peltier modules (Laird Electronics DA-011-05-02). The chambers with flexible polyimide heating strip as the heating element are controlled by GW INSTEK GPD3303D benchtop power supply. The chambers with peltier modules as the heating/cooling elements are controlled by Wavelength Electronics WTCP5V5A PWM Temperature Controller mounted on evaluation boards from the same company. The lighting setup is built with Thorlabs parts, and the cameras used are Basler cameras. For chambers with thermistors, 10 kΩ thermistors are attached to the bottom of wafers using TG-PP10-50 thermal putty, then covered by tape.

Behavioral data was pre-processed to get a trajectory using a custom MATLAB script where the organism’s position is determined at each frame by thresholding and size-filtering until the organism is isolated, then obtaining the centroid of the organism. The output is a time series of (x,y) position with the starting point set to (0,0), as well as the size (pixels) and perimeter at each frame. One caveat is that because trajectories are centroid trajectories, there can be some recorded displacements that are less related to motion along the trajectory and more related to organism shape changes.^62^ To mitigate the effects of the latter, for the windrose plots in Fig. 2 c-d, we filtered out data points where the organism centroid moved equal to or less than 0.0601mm (resolution limit) in both x and y.

### Orientation experiment

Similar to other behavior experiments, *T. adhaerens* were placed in nutrient-free ASW for 4-8 hours prior to behavioral experiments in order for them to shed their undigested algae coat so as to remove chemical information from algae to isolate the effect of thermal information.

For the orientation experiment, we used a Squid tracking microscope which was developed in our lab and described by Li et al.^33^ The organism was placed in a circular glass-bottom chamber (WillCo Wells, GWST-5040). Polystyrene “sticky microspheres” were prepared according to a procedure described by Prakash et al.^63^ Briefly, PolyLink Protein Coupling Kit (for COOH microparticles, Polysciences, Catalogue number 24350-1) was used to coat COOH - functionalized microspheres (31.7 *μ*m diameter, from Spherotech CM-300-10) with Wheat Germ Agglutinin (Molecular Probes, W11262). Then, the “sticky microspheres” were sprinkled onto the organisms using a micropipette. The organism was tracked under the tracking microscope until the stage reached its limit.

The outputs of the tracking program include a TIFF stack (acquired at 0.5fps) and the stage coordinates. We manually tracked the positions of 2 beads over the course of the experiment. Using Fiji we determined the angle *θ* as labeled in Fig.2f as the angle between the line drawn from bead 1 to bead 2 and the positive x-axis.

### Quantitative metrics

Evident in the raw trajectories in Fig. 1, the behavioral phenotypes that we observe show a lot of complexity across time scales. We thus chose to characterize the data by a myriad of metrics described below. Increasing the number of dimensions of each trajectory in the space of quantitative descriptors allows us to have a better toolkit that we can now use to explore possible mechanisms behind the phenotype by introducing perturbations into the assay that we have developed.

As measures of directedness of motion, we first considered mean velocities relative to thermal gradient axis. Mean velocities parallel and orthogonal to thermal gradient axis are determined as mean “momentary velocities” across sliding window length of *n* frames projected onto the thermal axis (or orthogonal axis), and the average is across 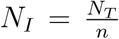 non-overlapping intervals (unless specified - for some metrics where we used bigger window sizes, we opted for overlapping windows). In our analysis we used non-overlapping sliding windows with *n* = 2 frames, and *δt* = *n*(30*s/frame*) = 1*min* unless otherwise indicated. Statistical tests were done using Scipy with Mann-Whitney U rank test.

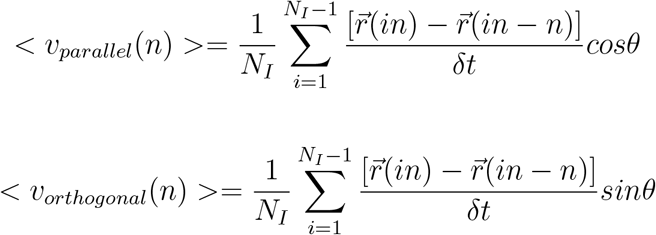

Mean squared displacement (MSD) is a common way to characterize the trajectories. We computed MSD using window size of 1 (30s intervals - highest temporal resolution available for our trajectories). Because we are interested in characterizing the directedness of motion along the thermal axis, we separated the MSD calculations into MSD parallel to the gradient axis (*MSD*_*parallel*_) and MSD orthogonal to the thermal axis (*MSD*_*orthogonal*_). The slope of the log-log MSD plot, also referred to as the *α* value, is a simple measure reflecting directional persistence; its value is 1 for random motion and 2 for a straight (ballistic) motion.^62^ *α <* 1 corresponds to subdiffusive motion, and 1 *< α <* 2 corresponds to superdiffusive motion.^64^

The straightness index (Eqn 4)^65^ is a simple ratio of end-to-end displacement (D) to total distance traveled (L).^66^

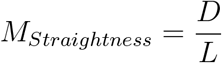

It is a good metric for quantifying the straightness of the trajectory but like MSD, is blind to direction. The directionality index (Eqn 4) is a slight modification of the straightness index to get an idea of efficiency of moving towards the thermal stimulus.

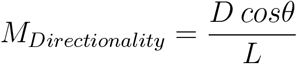

Path persistence length was determined as described in [^35^] using the equation:

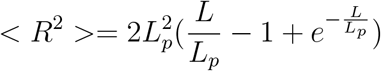

For incremental values of L (window length - time is not fixed here, only the total distance traveled in each window) up to 2cm, *< R*^2^ *>* was calculated as the averaged squared displacement between non-overlapping contours of length L. Then, the parameter *L*_*p*_ was determined using a least regression fit in MATLAB. Then, plotting *L* against *L*_*p*_, we determined the window length *L* for which we get the longest persistence length.

Another way to characterize of persistence in movement direction is to fit the root-meansquared-displacement (RMSD) curves using Taylor’s equation^67^ (Eqn 6):

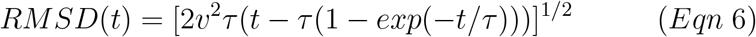

where the fit parameters *v* (mm min-1) is ballistic velocity and *τ* (min) is the decorrelation time-scale of the direction of movement.

For trials with thermal gradient, we noted that for most trials, the RMSD plot can actually be better fit with

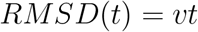

which is for ballistic motion. This is what we did for Fig.3f.

For MSD and RMSD calculations, in order to normalize for the trajectory length, and to avoid any artifacts that may be associated with stagnant periods near the beginning (e.g. some organisms take some time to unfold) and end (e.g. some organisms touch the edge of the arena) of some trials, we computed the MSD along thermal gradient axis using the 200 time points between [last frame-200-10, last frame-10], which corresponds to a 100 min time window near the end of the trajectory, ending 5 minutes before the end of each trial.

Complexity metrics such as DFA and Shannon entropy were computed using Neurokit. ^68^ Detrended fluctuation analysis (DFA) was originally introduced as a long term memory metric that is able to differentiate local patchiness (mosaic structure of DNA where there are lots of local excess of a particular base pair) from long-range correlations.^69^ This is useful for our data because we also see “local patchiness” in momentary velocities, as the trajectories are “jittery”, with segments where the centroid of the organism is mostly stationary.

Lévy distributions ^36^ have densities:

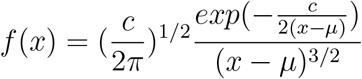

To determine parameters *c* and *μ* for our observed step length distributions, we combined the step length data (we used overlapping window sizes of 10 minutes and used the portion of the step length parallel to the gradient axis) from all trials for each condition (thermal gradient and control), then used least squares regression fitting in MATLAB.

### Drug perturbation assays

Drug perturbation assays are done in the same chamber type as behavioral assays (2.06 inch ID PVC tubing on silicon wafer with flexible heating strip at bottom of wafer), and organisms are similarly starved prior to experiment. Naringenin (TCI chemicals, product number N0072) was dissolved in DMSO (Sigma-Aldrich, CAS number 67-68-5) to make a 10 mM stock solution. A new stock solution is made fresh on the day of each experiment. This stock solution is then diluted in ASW to a 150mL solution immediately prior to addition, so that when added to the 25mL chamber, the final naringenin concentration would be 40 *μ*M (we did a concentration screen and picked the lowest concentration at which we saw a behavioral effect). Similarly, Flufenamic acid (Sigma-Aldrich, CAS number 530-78-9) was dissolved in DMSO (Sigma-Aldrich, CAS number 67-68-5) to make a 10 mg/mL stock solution. This stock solution is then diluted in ASW to a 150mL solution, so that when added to the 25mL chamber, the final FFA concentration would be 10 *μ*M (we did a concentration screen and picked the highest concentration that does not kill the organism).

## Supporting information

Supplementary Video 1: T. adhaerens thermotaxis behavior

Supplementary Video 2: Thermotaxis under shallow temperature gradient

Supplementary Video 3: Inhibition of thermotaxis by naringenin

Supplementary Video 4: Response to sudden change in gradient direction

Supplementary Video 5: Organism with anomalous perimeter-area ration has difficulty performing thermotaxis

Supplementary Video 6: Thermotaxis under "confounding" flow

Supplementary Video 7: T. adhaerens doesn't have fixed orientation while performing thermotaxis

## Acknowledgements

We thank Chew Chai, Pranav Vyas, and Clarice D. Aiello for engagement during conception phase of the project. We thank Hongquan Li and Deepak Krishnamurthy for help with Squid microscopy setup. We thank Vipul Vachharajani, Raphael Eguchi and all members of the Prakash lab for discussions and useful feedback. GZ gratefully acknowledges the support of the Natural Sciences and Engineering Research Council of Canada (NSERC) PGS-D program as well as support by Stanford University Agilent Fellows Program. This work is supported by HHMI Faculty fellowship (MP), BioHub Investigator Fellowship (MP), Schmidt Innovation Fellowship (MP), Keck Foundation Research Grant and NSF CCC (DBI1548297) grant.

## Notes

### Competing Interest Statement

The authors have declared no competing interest.

