## Supplementary Information for "Thermotaxis in an apolar, non-neuronal animal"

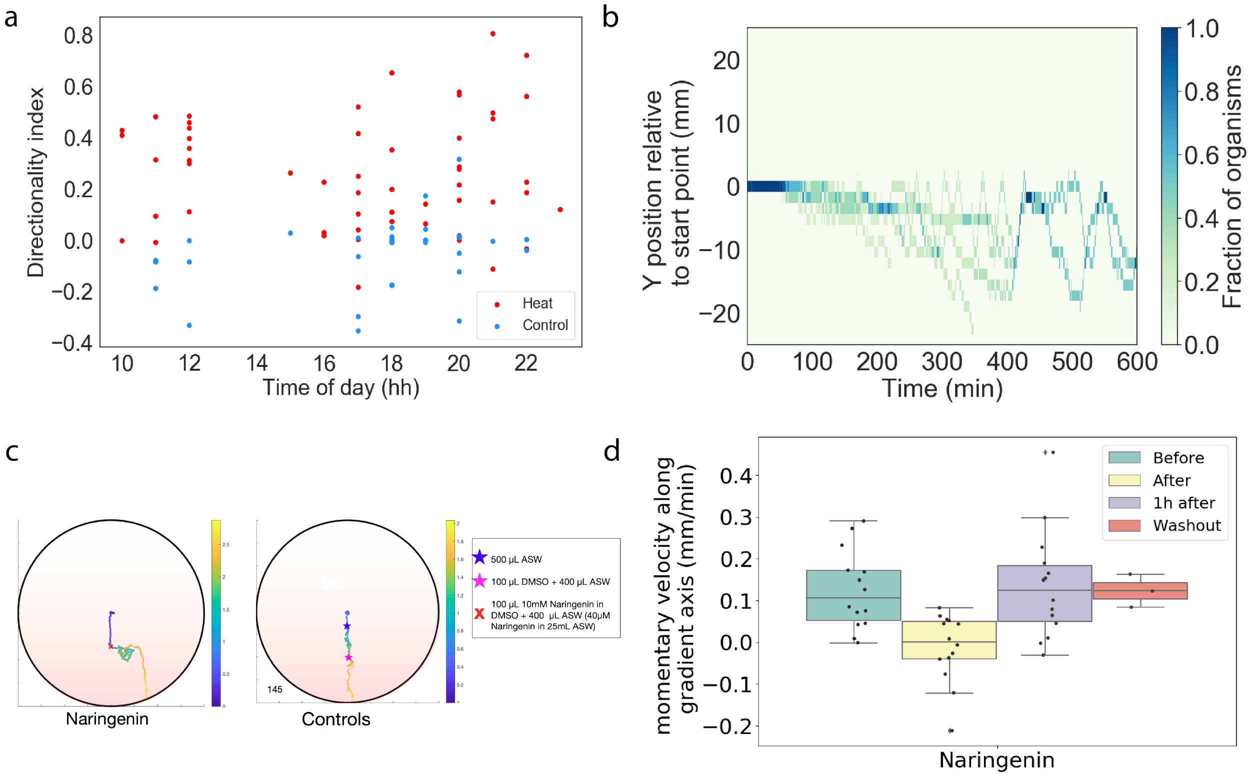


Figure 1: Additional control experiments. (a) There is no apparent circadian dependence of directionality index in *T. adhaerens* thermotaxis. Regardless of hour of day when the experiment is started, the directionality index of trials with a thermal gradient tends to fall above 0 (indicating positive thermotaxis). (b) To ensure that the organism is sensing temperature gradient, we added a stir bar to generate an external, strong 3mm/s confounding flow, but despite the flow, this kymograph shows that the organism still appears to be able to find the thermal gradient, venturing towards the warmer side of the arena within 600 minutes of the start of the experiment. (c-d) More details on control experiments for thermotaxis inhibition by naringenin. (c) Representative trajectories showing naringenin addition causes a sharp, reversible disruption of thermotaxis (left), but that this disruption is not due to the DMSO and ASW that was added with the naringenin (right). For both representative trials, the heat source is located at y=-25mm. (d) Boxplot showing $V_{parallel}$ in the 30 minute time windows starting at indicated time points relative to Naringenin addition. For washout trials (red), Naringenin is washed out by serial dilution approximately half an hour after addition. Recovery of thermotaxis behavior occurs naturally after an hour of drug addition or following washout of drug.

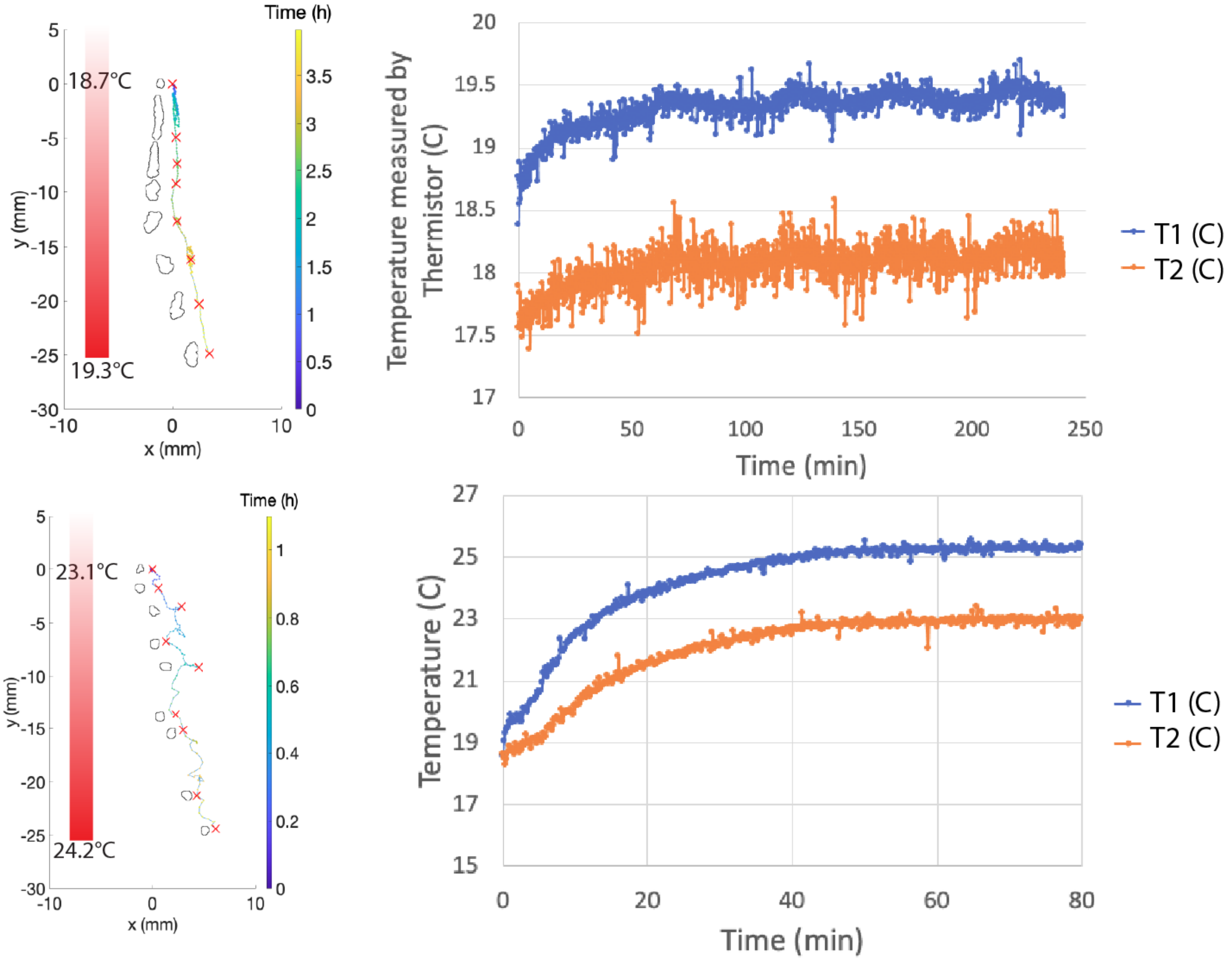


Figure 2: Temperature profiles for sample trajectories shown in main text Fig.1. Temperature readout is from thermistors, T1 is the temperature at y=-25mm, at the warmer side of the arena; T2 is the temperature at y = 25mm, at the cooler side of the arena. These are two examples of thermotaxis assays at different absolute temperatures for T1, and clearly positive thermotaxis occurs in both cases.

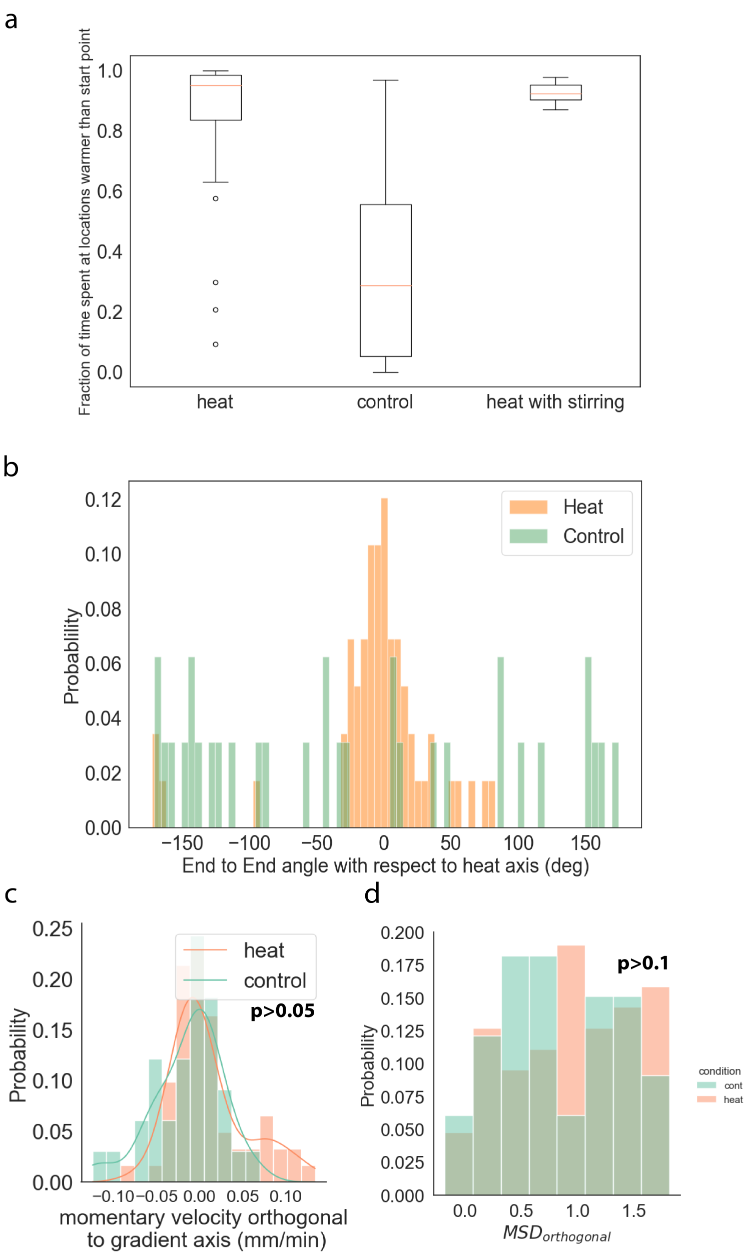


Figure 3: Additional quantitative metrics for understanding motility trajectories. (a) When a thermal gradient is present, the organism spends more time exploring locations that are warmer than the starting position. This holds even when an external water flow is generated in addition to presence of thermal gradient. (b) Distribution of end-to-end angle of trajectory (with respect to thermal gradient axis) for experiments with (orange) and without (green) thermal gradient. When gradient is applied, end-to-end trajectories are highly aligned with the thermal axis. The distribution for trials with thermal gradient has a mean of 0.87 degrees and standard deviation of 50 degrees. (c-d) Orthogonal components of momentary velocity (c) and slope of log-log MSD (d) are not significantly affected by application of thermal gradient. Sliding window size is 1min, non-overlapping.

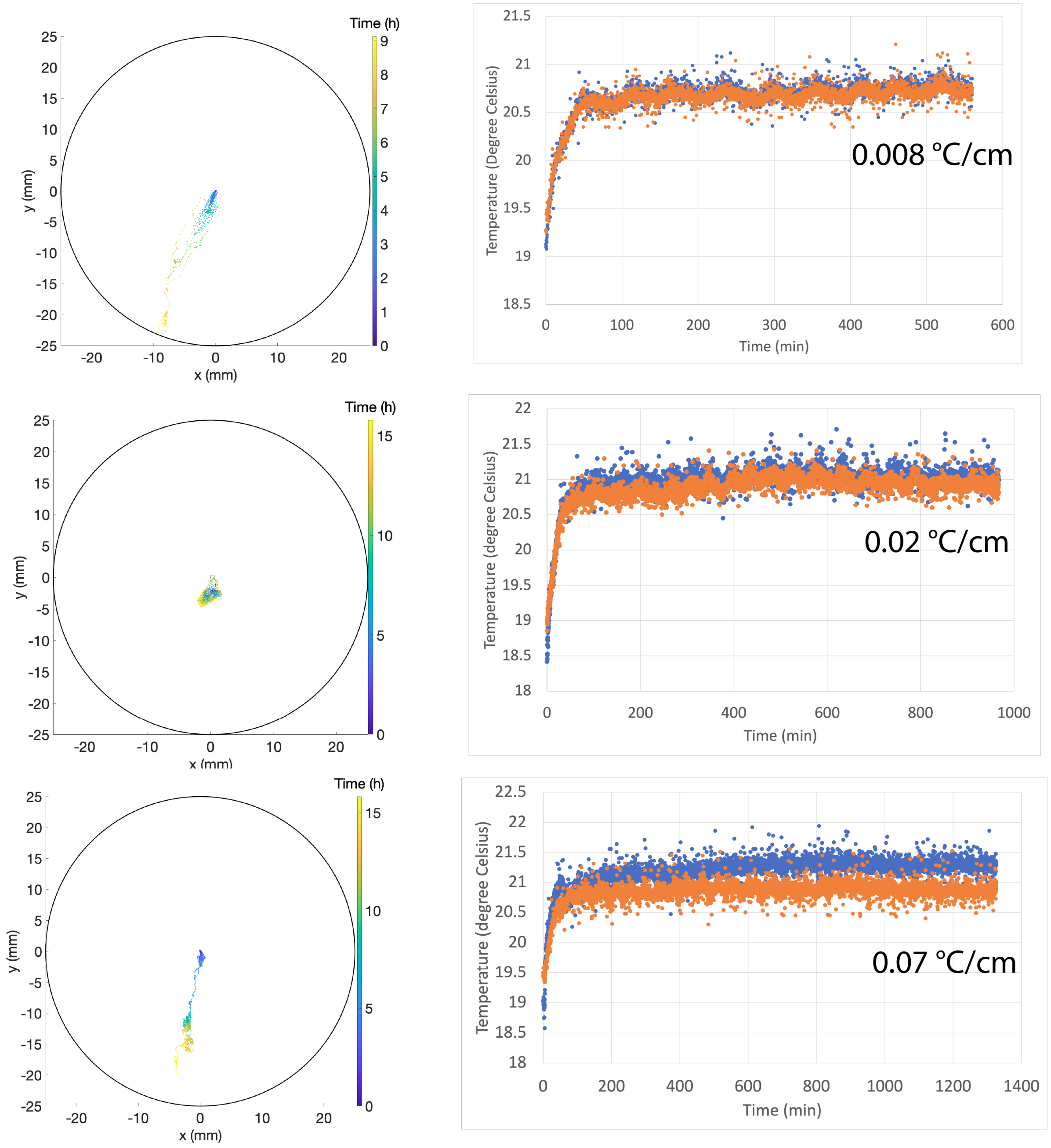


Figure 4: Single organism trajectories (left) and temperature profiles (right) for three trials with the lowest thermal gradients presented in this work. Two out of three organisms were able to find and move towards the gradient, and the organism presented in the topmost trajectory (Supplementary Video 2) was able to navigate the shallowest gradient we generated (0.008 °C/cm)

Table 1: Hypothetical TRP channel homologs have been previously reported in *T. adhaerens* genome. Within the reported channels, TRPV1-2, TRPM2-4 have previously been reported to show thermal activation. Percent similarity to *Mus musculus* TRP sequences are also shown.

| Hypothetical *T. Adhaerens* TRP channel homologs (protein database accession) | % Similarity with *Mus musculus* | Reported temperature sensitivity (mammalian) |
|---|---|---|
| TRPM1 (XP_002115051) | 23.3% | |
| TRPM2 (XP_002114984) | 23.2% | ✓ |
| TRPM3 (XP_002114617) | 32.7% | ✓ |
| TRPM4 (XP_002116621) | 32.7% | ✓ |
| TRPML (XP_002113929) | 54.3% | |
| TRPP1 (XP_002115395) | 56.8% | |
| TRPP2 (XP_002110052) | 22.0% | |
| TRPV1 (XP_002114713) | 23.6% | ✓ |
| TRPV2 (XP_002118208) | 25.8% | ✓ |
